# Delayed onset and heterogeneous collective organization characterize twitching motility in *Acinetobacter baumannii*

**DOI:** 10.64898/2026.07.01.735841

**Authors:** Clara Dessenne, Anaïs Henriques, Adeline Courseaux, Corentin Spriet, David Dauvillée, Olivier Vidal, Yannick Rossez

## Abstract

Type IV pili (T4P) mediate twitching motility and contribute to surface colonization, biofilm formation, and host interactions in *Acinetobacter baumannii*. However, the prevalence, dynamics, and diversity of twitching motility across *A. baumannii* populations remain poorly understood. Here, we compared twitching motility in a collection of 35 *A. baumannii* strains originating from clinical, environmental, and animal sources, using *Pseudomonas aeruginosa* PAO1 as a reference. Standardization of assay conditions revealed a strong influence of agar composition on twitching motility, with Eiken agar supporting the most robust surface translocation. Under these conditions, 14 of 35 *A. baumannii* isolates exhibited detectable twitching motility. Time-lapse microscopy revealed major differences between *A. baumannii* and *P. aeruginosa*. Whereas PAO1 initiated twitching within minutes after inoculation and formed characteristic multicellular rafts, motile *A. baumannii* strains displayed a prolonged non-motile phase before movement initiation and exhibited distinct patterns of collective organization. Two major expansion phenotypes were identified, termed Homogeneous Front (HF) and Raft-Like Front (RLF), together with Early-Onset Motility (EOM) and Delayed-Onset Motility (DOM) subgroups. Quantitative analyses further revealed substantial variation in speed, directional persistence, and migration dynamics among strains. Because a majority of isolates were non-motile, we investigated the contribution of the minor pilin FimT. Although deletion of *fimT* abolished twitching motility and specific substitutions modulated motility efficiency, sequence variation in FimT alone could not account for the observed phenotypic diversity. Collectively, these findings reveal extensive heterogeneity in T4P-mediated surface motility in *A. baumannii* and identify delayed twitching activation and distinct collective migration strategies as key features of surface colonization in this species.

**Highlights:**

- Population analysis reveals extensive twitching diversity in *A. baumannii*
- Delayed motility onset distinguishes *A. baumannii* from *P. aeruginosa*
- Homogeneous and raft-like fronts define distinct migration strategies
- FimT contributes to motility but not all non-motile phenotypes

## Introduction

The earliest description of what is now known as twitching motility, originally termed gliding, dates back to Hans Lautrop’s work in 1961 on *Bacterium anitratum* (Lautrop, 1961). Lautrop characterized this surface-associated movement as “slow, hesitant and intermittent,” drawing parallels with the motility observed in other Myxobacterales, including *Moraxella/Cytophaga lwoffii* (later renamed *Acinetobacter lwoffii*). Based on these motility features on solid surfaces, together with cellular morphology and the absence of microcyst formation, Lautrop proposed reclassifying *Bacterium anitratum* as *Cytophaga anitrata*. Concurrently, other authors referred to this organism as *Acinetobacter anitratum*, *Acinetobacter calcoaceticus* subsp. *anitratus*, or *Acinetobacter calcoaceticus* (Baumann et al., 1968; Thornley, 1967). In subsequent years, the bacterial strain in which this phenomenon was first observed was reclassified as *Acinetobacter baumannii* (Bouvet and Grimont, 1986). Earlier, Lautrop (1965) had coined the term “twitching motility”, providing a unifying framework for this mode of surface translocation that was later recognized in *A. baumannii*. Despite this historical primacy, detailed microscopic and mechanistic descriptions of twitching motility in *A. baumannii* remain remarkably scarce, especially when compared to the extensive body of work available for *Pseudomonas aeruginosa* (Burrows, 2012; Henrichsen and Blom, 1975; Kühn et al., 2021; Lautrop, 1961; Mattick, 2002; Meacock et al., 2021; Semmler et al., 1999) Interest in *A. baumannii* motility has recently intensified due to its emergence as a major human pathogen and its increasing resistance to multiple classes of antibiotics, creating an urgent need for new therapeutic strategies (Miller and Arias, 2024). In this context, targeting bacterial motility, an essential determinant of surface colonization, biofilm formation, and host interaction, represents a promising anti-virulence approach, offering the possibility to disarm the pathogen and limit its spread without directly exerting selective pressure on bacterial viability (Sun et al., 2026). Twitching motility, mediated by type IV pili (T4P), is not restricted to *A. baumannii* but is shared by a wide range of bacteria, including *P. aeruginosa*, *Vibrio cholerae*, *Caulobacter crescentus*, *Myxococcus xanthus*, *Neisseria* spp., and even some Gram-positive species such as *Clostridium* spp., and *Streptococcus sanguinis* (Berry et al., 2019; Craig et al., 2019; Mattick, 2002; Piepenbrink and Sundberg, 2016; Varga et al., 2006). To date, three subtypes of T4P have been described (Type IVa, Type IVb, and Type IVc) with T4aP being the most prevalent and best characterized. T4aP are widely distributed among Gram-negative bacteria including *A. baumannii* and *P. aeruginosa* (Ellison et al., 2022). This motility mechanism is critical for colonization of hydrated environments and is strongly influenced by substrate physicochemical properties (Carabelli et al., 2020; Gomez et al., 2023). Twitching motility relies on the coordinated, multicellular activity driven by cycles of T4aP extension, surface attachment, and retraction, enabling bacteria to efficiently explore and colonize surfaces. Beyond motility itself, T4aP plays a central role in the establishment and spatial organization of bacterial communities, contributing to early surface colonization, biofilm formation, and the development of complex multicellular structures such as *Myxococcus* fruiting bodies (Bonner et al., 2006; Klausen et al., 2003; Yu et al., 2025). These collective behaviors are key determinants of bacterial persistence in both environmental and host-associated settings. T4aP functions also directly contribute to host colonization and pathogenicity by promoting adhesion, surface persistence, and tissue interaction during infection (Ahmad et al., 2023; Zolfaghar et al., 2003). In *A. baumannii*, transcriptional analyses have shown that T4aP-associated genes are upregulated during growth in human serum, suggesting an active role for pili-mediated functions during bloodstream infection and host adaptation (Jacobs et al., 2012; Murray et al., 2017). Furthermore, recent studies have shown that virulence of *A. baumannii* may depend on the pilus tip adhesin ComC (also known as PilY), underscoring the critical role of T4aP-mediated adhesion in host interaction and infection (Iruegas et al., 2023). In the same study, attention was also given to the minor pilin FimT. FimT contains a GspH Pfam domain, a feature typically associated with pseudopilins involved in bacterial Type II secretion systems (T2SS) (Naskar et al., 2021). This domain architecture suggests potential functional convergence between T4aP minor pilins and pseudopilin components of secretion systems, further supporting the idea of structural and functional diversification within the T4aP machinery.

Despite the recognized importance of T4aP in *A. baumannii*, fundamental questions remain regarding how twitching motility is deployed at the microscopic scale, whether different strains exhibit distinct patterns of surface-associated movement and collective organization, and to what extent natural variation in T4P-associated proteins contributes to this phenotypic diversity. Recent comparative genomic analyses identified lineage-specific diversification of several T4aP components, including ComC/PilY1, FimU, PilV, and the minor pilin FimT, suggesting that natural variation within the T4aP machinery may contribute to functional diversification (Iruegas et al., 2023). However, the relationship between this genetic diversity and the diversity of twitching phenotypes remains largely unexplored. We therefore compared twitching motility across a collection of 35 A. baumannii strains representing environmental, animal, and clinical isolates (Dessenne et al., 2024; Wilharm et al., 2026), together with the clinically relevant strain AB5075 (Jacobs et al., 2014), using *P. aeruginosa* PAO1 as a reference under standardized experimental conditions (Stover et al., 2000). Combining quantitative motility assays with time-lapse microscopy allowed us to document previously unrecognized diversity in the temporal and collective behaviors associated with T4aP-mediated surface translocation. Finally, we investigated whether natural sequence variation in the minor pilin FimT contributes to this phenotypic diversity through complementary genetic and structural analyses.

## Methods

### Strains and growth conditions

Bacteria were cultivated in lysogeny broth (LB)-Lennox (for 1 liter 10g of tryptone, 5g of yeast extract and 5g of NaCl) supplemented with appropriate antibiotics (ampicillin 100 µg/mL; apramycin 50 µg/mL for *E. coli* and 200 µg/mL for *A. baumannii*) or 37°C at 180 rpm.

### Genetic engineering of *Acinetobacter baumannii*

The Δ*fimT* (ABUW_0648) deletion mutant was generated through a standard allelic exchange method using the counter selectable suicide vector pGP704-Sac-Apr conferring apramycin resistance to *Acinetobacter baumannii*. This plasmid was obtained by replacing the *Bla* resistance gene of pGP704-Sac28 (Metzger et al., 2019) by the *aac(3)-IV* open reading frame from pBECAB-Apr (Wang et al., 2019) by Gibson assembly (NEB, Biolabs) using the following couples of primers *pGP704-ApR_fwd/rev and ApR_fwd/rev* for pGP704 and *aac(3)-IV* ORF amplification respectively. The integrity of the final construct was checked by Sanger sequencing (Eurofins Genomics, Ebersberg, Germany). The pGP704-Sac-Apr-Δ*fimT* deletion construct was obtained by joining 1kb of the upstream and 1kb of the downstream regions flanking the *fimT* locus by overlap extension PCR using Q5 DNA polymerase. The 2 kb PCR product was then cloned in the unique *Xba*I restriction site in pGP704-Sac-Apr. The integrity of the final construct was checked by Sanger sequencing (Eurofins Genomics, Ebersberg, Germany). The deletion construct was transferred to *A. baumannii* through mating with *E. coli* S17-1lpir. Individual *A. baumannii* apramycin resistant clones were cultivated overnight without antibiotics before being spread onto LB plates containing 10% sucrose. Apramycin sensitive colonies carrying either the WT or the deleted *fimT* copies were identified by PCR using primers surrounding the gene. The broad host-range and multicopy pASG-1 plasmid (Godeux et al., 2018) carrying the super folder GFP gene under a strong constitutive promoter was used as a backbone for complementation. The pASG-*fimT* plasmid was obtained by replacing the *Nde*I/*Pst*I fragment carrying SFGFP by the *fimT* ORF which was amplified by PCR with primers allowing the introduction of both a *Nde*I (5’ end) and a *Pst*I (3’ end) restriction enzyme sites.

### Macroscopic motility

Petri dishes containing 1% agar in LB were inoculated with 5 µL of bacterial suspension in exponential phase (OD₆₀₀ = 0.4). Plates were then incubated at 37 °C for 72 h and sealed with parafilm to prevent agar desiccation throughout the incubation period. Different experimental trials were performed using modified LB media in combination with several agar sources to evaluate potential effects of agar composition on bacterial behavior. The agar types tested included BD Difco™ Granulated Agar (product code 11793523), Thermo Scientific™ Agar Powder (CAS 9002-18-0), Thermo Scientific™ Oxoid Agar (LP0011B), MicroAgar (Duchefa Biochemie, CAS 9002-18-0), and Eiken agar (Biotechnik Gerbu, E-MJ00). Agar media were prepared according to manufacturers’ instructions at identical final concentrations to ensure comparability across experiments. For reproducibility, Petri dishes were poured with a standardized volume of 20 mL of medium per plate. All plates were prepared extemporaneously immediately prior to use to minimize variations in agar hydration and surface properties that could influence motility behavior. FimT-dependent twitching motility was quantified as the expansion halo area on agar plates. Residual colony expansion was consistently observed under non-twitching conditions and was measured as background, which was subtracted from all measurements. Data were normalized to the WT (WT = 100%). As a result, strains lacking detectable twitching motility could yield negative values due to background correction. For visualization only, values below zero were rescaled using a piecewise linear transformation, allowing non-motile strains to be displayed on a 0–10% scale while preserving the relative differences among twitching-positive strains.

### Microscopic motility

µ-Dish 35 mm glass-bottom dishes (Ibidi, FluoroDish FD35-100) were filled with 3 mL of LB medium supplemented with 1% Eiken agar and allowed to solidify prior to inoculation. Each dish was inoculated with 4 µL of bacterial suspension prepared in exponential growth phase. All dishes were sealed with parafilm to prevent medium desiccation during incubation and imaging. Dishes were incubated and directly monitored at 37 °C using a Nikon Biostation IM-Q live-cell imaging system. Bacterial behavior was recorded at multiple time points between 0 and 10 h post-inoculation. Time-lapse videos were acquired by capturing images every 2 s over 3 min intervals using 40× and 80× objectives. Angular velocity (rad·s⁻¹) was calculated as the rate of change of the trajectory angle over time and used as a measure of directional persistence during surface translocation.

### Protein structure prediction and analysis

The three-dimensional structures of FimT variants (AB5075, ABGW 5, ABGW 6, and ABGW 27) were predicted using AlphaFold (Abramson et al., 2024). Structural superpositions and alignments with the WT FimT reference structure from strain AB5075 were performed using UCSF ChimeraX (version 1.11.1) (Meng et al., 2023). Local and global structural metrics, including root-mean-square deviation (RMSD) values, Cα atomic distances, and solvent-accessible surface area (SASA), were calculated within the ChimeraX environment. Electrostatic potential variations at the molecular surface were visualized and evaluated through Coulombic surface coloring analysis.

### Video analysis

Time-lapse videos were analyzed using Fiji (ImageJ distribution) (Schindelin et al., 2012). Image sequences were first preprocessed by adjusting contrast and applying intensity thresholding to enhance bacterial detection. A binary mask was then generated to improve object segmentation and reduce background noise prior to tracking analysis. Single-cell tracking was performed using the TrackMate plugin. Spot detection was conducted using the Laplacian of Gaussian (LoG) detector, with parameters optimized for bacterial cell size and signal-to-noise ratio in each dataset. Tracking was subsequently performed using the Linear Assignment Problem (LAP) tracker, allowing gap closing and linking of trajectories across frames when appropriate. Tracking outputs were visually inspected to verify detection accuracy and correct trajectory reconstruction. Quantitative tracking data, including positional coordinates and trajectory statistics, were exported in spreadsheet format for downstream analysis of bacterial motility parameters. For figures 2A, 2B, 10A and 10B, cell trajectories were reconstructed from 3-min time-lapse recordings by tracking bacterial x,y coordinates. For visualization, trajectories were temporally downsampled by plotting cell positions every 30 frames (60 s; t = 0, 1, and 2 min), providing a simplified representation of movement while preserving the overall trajectory.

### Statistical analysis

Statistical analyses were performed using GraphPad Prism version 8.0 (GraphPad Software, San Diego, CA, USA). Depending on the dataset, statistical significance was assessed using either one-way analysis of variance (ANOVA) followed by Tukey’s multiple comparisons post hoc test or the Kruskal-Wallis test. Data represent independent biological replicates. Differences were considered statistically significant at *p* ≤ 0.05.

## Results

### Agar composition influences twitching motility and species-specific behaviors

To establish reproducible conditions for twitching motility analyses, five agar formulations were compared using *P. aeruginosa* PAO1 and *A. baumannii* AB5075. Eiken agar supported the most pronounced twitching motility in both species, whereas Thermo agar promoted robust motility in PAO1 but only limited movement in AB5075, indicating species-specific responses to substrate composition (Figure 1). Using these standardized conditions, twitching motility of PAO1 and AB5075 was compared by time-lapse microscopy. As previously described, PAO1 rapidly formed multicellular rafts that migrated collectively away from the inoculation site (Figure 2A; Movie 1), generating densely packed coordinated groups with occasional finger-like projections (Semmler et al., 1999; Turnbull and Whitchurch, 2014). In contrast, AB5075 displayed a more homogeneous radial expansion, with cells dispersing outward in all directions from the inoculation site (Figure 2B; Movie 2). Quantitative tracking over 480 min revealed marked differences in twitching dynamics (Figure 3). Across the full dataset, PAO1 exhibited higher translational speeds than AB5075, with a mean speed of 0.322 ± 0.120 µm·s⁻¹ compared with 0.168 ± 0.095 µm·s⁻¹ for AB5075 (Figure 3A). Maximum speeds reached 0.583 ± 0.169 µm·s⁻¹ for PAO1 and 0.417 ± 0.197 µm·s⁻¹ for AB5075 (Figure 3B), whereas AB5075 displayed a slightly higher angular velocity (0.817 ± 0.200 rad·s⁻¹) than PAO1 (0.729 ± 0.214 rad·s⁻¹), consistent with more frequent reorientation events (Figure 3C). Time-resolved analyses further highlighted differences in twitching initiation and progression (Figures 3D and E). PAO1 initiated twitching approximately 15 min after inoculation and progressively increased its speed before reaching a plateau between 180 and 240 min, followed by a decline after 300 min (Figure 3F). The highest maximum speed recorded for PAO1 was 0.616 ± 0.231 µm·s⁻¹ during the 300-360 min interval (Figure 3G). Angular velocity peaked during the initial phase corresponding to twitching onset and remained stable thereafter (Figure 3H). In contrast, AB5075 exhibited a prolonged non-motile phase and initiated twitching after approximately 120 min. Speeds increased until a plateau was reached between 240 and 300 min, followed by a decline between 360 and 420 min and subsequent stabilization (Figure 3I). The highest maximum speed recorded for AB5075 was 0.567 ± 0.158 µm·s⁻¹ during the 360-420 min interval (Figure 3J). Angular velocity remained relatively constant throughout the motile phase (Figure 3K).

**Figure 1:**
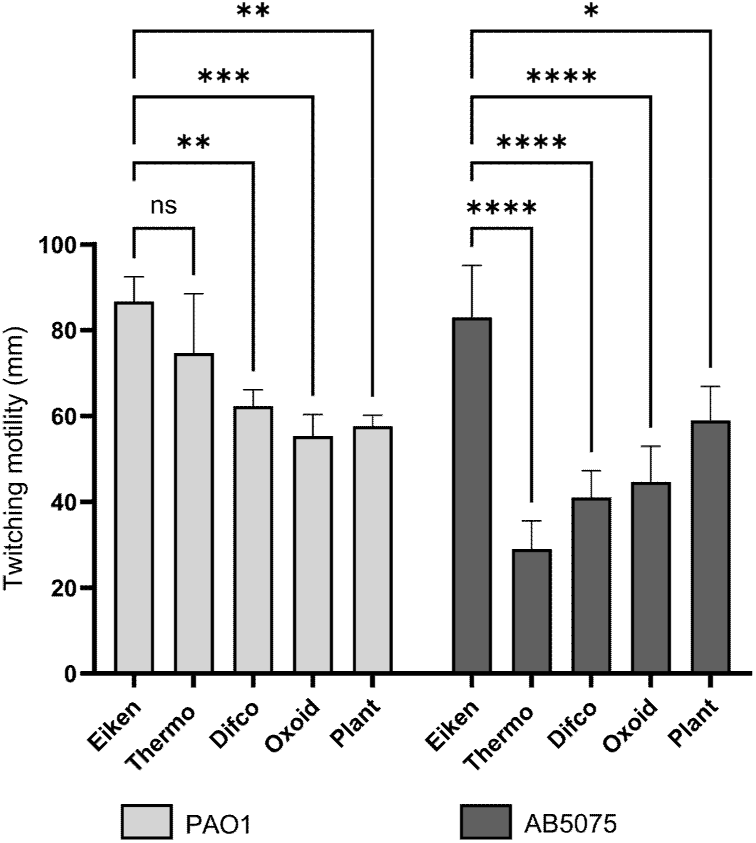
Effect of agar type on the twitching motility of *P. aeruginosa* PAO1 and *A. baumannii* AB5075. Representation of twitching motility measured in millimeters (mm) after 72h for the two bacterial species on different types of agar (Eiken, Thermo, Difco, Oxoid, Plant). Light gray bars represent the results for PAO1, while dark gray bars show those for AB5075. These variations suggest an influence of agar type on the motility capacities of the two species, potentially linked to differences in the composition or structure of the media. Statistical significances were determined by a two way ANOVA. (****, *p* ≤ 0.0001; ***, *p* ≤ 0.001; **, *p* ≤0.01).

**Figure 2:**
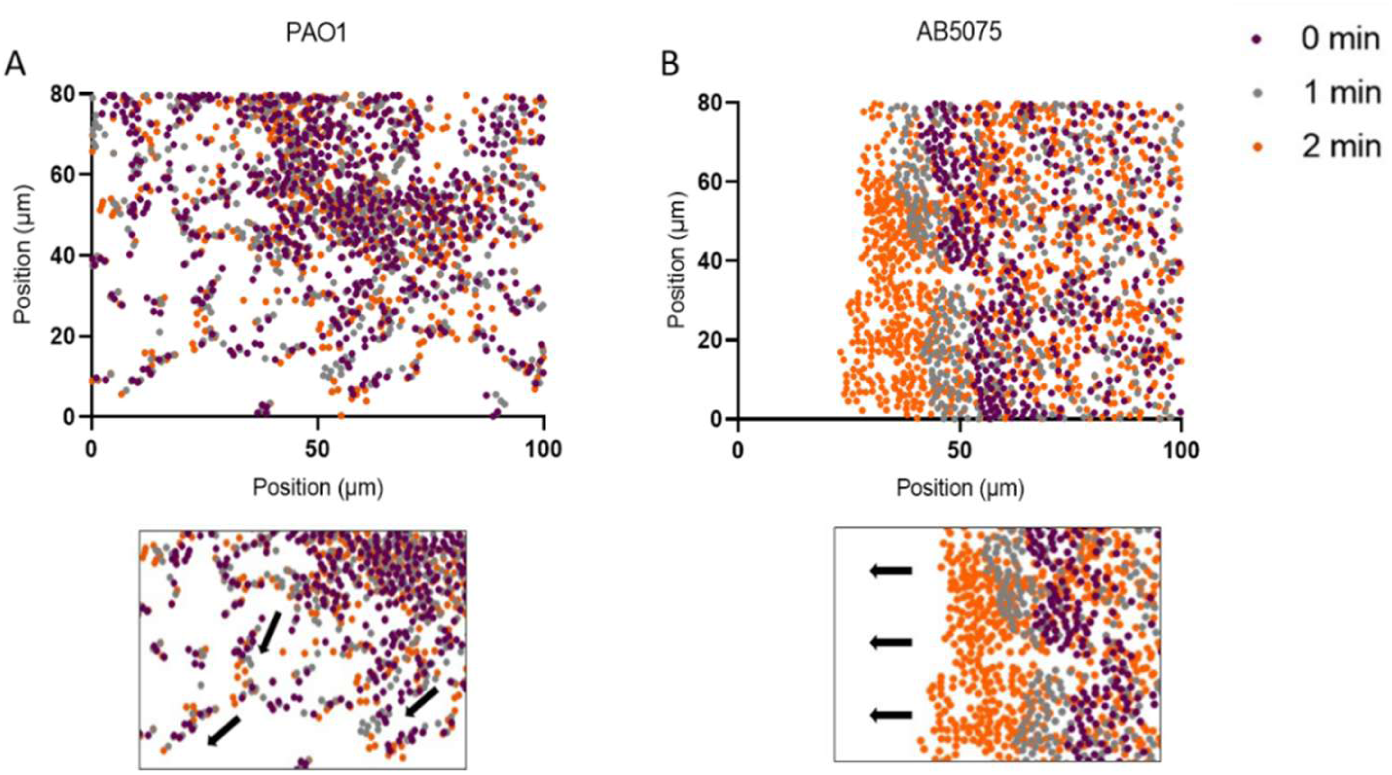
**Visualization of bacterial movement over time of *P.aeruginosa* PAO1 and *A. baumannii* AB5075**. Distinct modes of twitching motility in response to the same experimental conditions. T = 0 min purple points, T = 1 min gray points, and T = 2 min orange points at 3 hours. (A) Schematic representation of PAO1 movement during 2 min. The points represent the positions of bacteria on a surface expressed in µm. This strain shows an organization into migration arms, visible in the zoomed-in panel. Black arrows indicate the main axes of arm formation. (B) Movement of AB5075 bacteria at the same time points. The points also represent the positions of bacteria on a surface expressed in µm. This strain forms a homogeneous migration front visible across the population. Black arrows indicate the general direction of the front’s migration.

**Figure 3.**
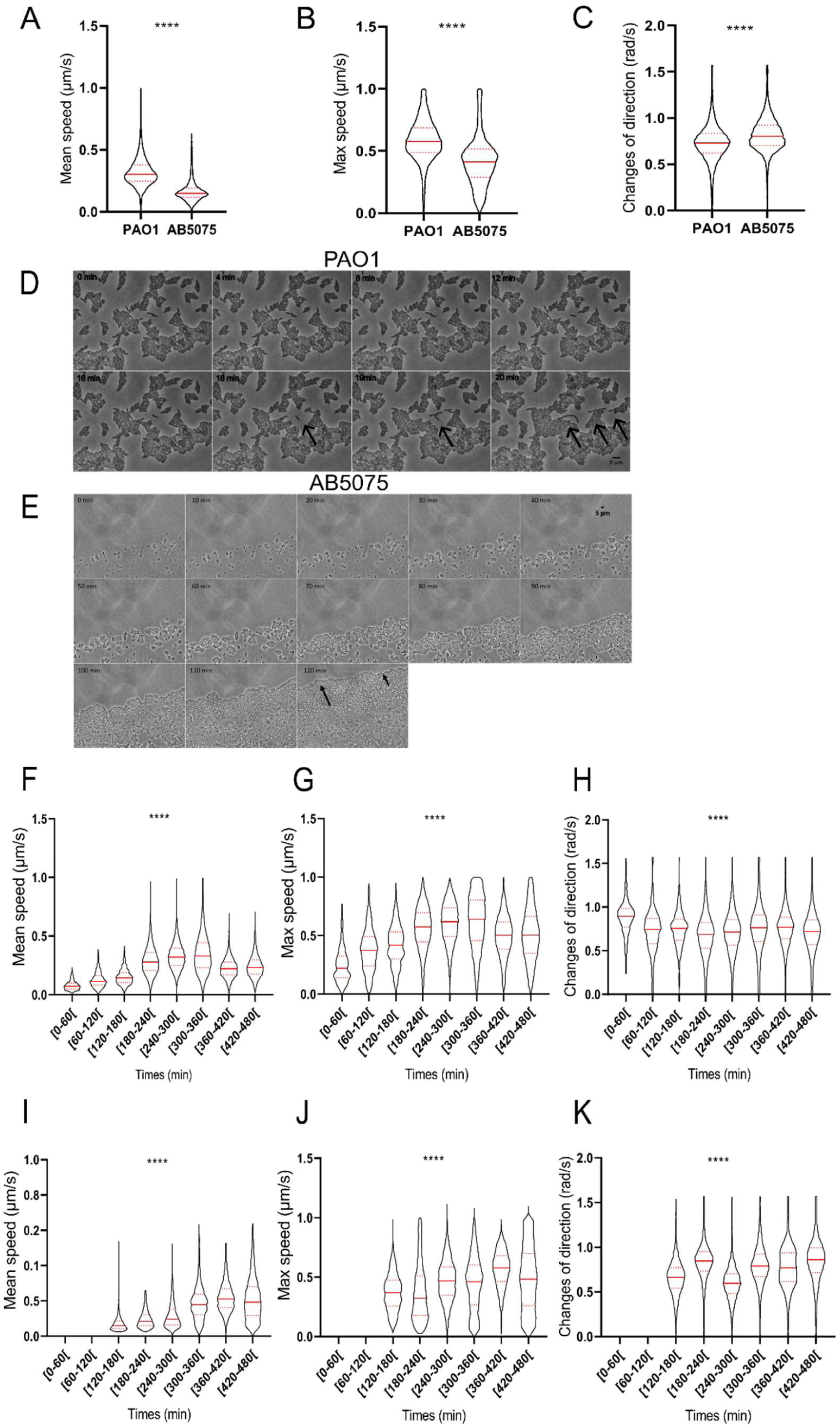
Comparative microscopic analysis of twitching motility dynamics in *P. aeruginosa* PAO1 and *A. baumannii* AB5075. Average motility characteristics of *P. aeruginosa* PAO1 (n = 20,797 cells) and *A. baumannii* AB5075 (n = 12,554 cells) were analyzed from time-lapse microscopy videos recorded over 8 h. (A) Mean speed (µm/s), (B) maximum speed (µm/s), and (C) rate of directional changes (rad/s) calculated for each species. Differences in the kinetics of twitching motility initiation are illustrated in (D) and (E). Time-lapse microscopy images acquired at 40× magnification show the onset of twitching motility within the same field of view. (D) For *P. aeruginosa* PAO1, images were acquired every 5 min. (E) For *A. baumannii* AB5075, images were acquired every 10 min. From 0 to 110 min, the increase in cell density primarily reflects bacterial growth and cell division, with no detectable displacement. At 120 min, twitching motility becomes clearly apparent, marked by the dispersal of cells ahead of the population edge, followed by the initiation and progression of a coherent migration front. Temporal evolution of motility parameters is shown for *P. aeruginosa* PAO1 in (F-H) and *A. baumannii* AB5075 in (I-K). (F, I) Mean speed (µm/s), (G, J) maximum speed (µm/s), and (H, K) rate of directional changes (rad/s) plotted as a function of time. For *P. aeruginosa*, measurements were performed from 20 min to 8 h after inoculation, whereas for *A. baumannii*, measurements start at 2 h because of an initial latency phase during which the bacteria were non-motile. Each data point represents the average value of the tracked bacterial population at the corresponding time point. Statistical significance was determined using the Kruskal-Wallis test following two-way analysis of variance for panels A-C and one-way analysis of variance for panels F-K (****, *p* ≤ 0.0001).

### Diversity of twitching motility among *A. baumannii* strains

To determine whether the twitching behavior observed in AB5075 was representative of the species, a collection of 35 *A. baumannii* strains was screened for macroscopic twitching motility. Fourteen isolates displayed detectable motility, whereas 21 strains remained non-motile under the experimental conditions used (Figure 4). The motile isolates were subsequently selected for detailed microscopic analyses. Although most sequenced *A. baumannii* strains encode T4P-associated genes, not all exhibit functional motility *in vitro* (Al-Shamiri et al., 2021; Eijkelkamp et al., 2011), indicating that gene presence alone is insufficient to ensure motility. Microscopic analyses of the 14 motile strains performed two hours after inoculation revealed substantial variation in twitching behavior. Mean speeds varied among isolates, with ABVal4 and ABGW2, ABGW5, ABGW7, ABGW17, and ABGW23 displaying values comparable to AB5075, whereas ABVal1 and ABGW1, ABGW8, ABGW11, ABGW14, ABGW16, ABGW21, and ABGW30 exhibited higher velocities (Figure 5A). Maximum speeds were generally similar, although ABGW2 was slower than AB5075, while ABVal1 and ABGW8 displayed higher peak velocities (Figure 5B). Angular velocity remained relatively conserved among strains, with the exception of ABVal4, which showed reduced directional changes, and ABGW2 and ABGW5, which exhibited increased reorientation (Figure 5C). Analysis of colony-front organization identified two major patterns of collective behavior. Homogeneous Front (HF) strains displayed smooth radial expansion similar to AB5075 and included ABGW8, ABGW11, and ABGW21 (Figure 5D, movie 3 for ABGW8). In contrast, Raft-Like Front (RLF) strains formed patchy, loosely aggregated clusters at the leading edge and included ABGW1, ABGW2, ABGW7, ABGW14, ABGW16, ABGW17, ABGW23, ABGW30, ABVal1, and ABVal4 (Figure 5E, movie 4 of ABGW7). Twitching onset also varied among isolates. Four strains (ABGW1, ABGW8, ABGW11, and ABVal4) exhibited Early-Onset Motility (EOM) within approximately one hour, whereas the remaining strains displayed Delayed-Onset Motility (DOM), initiating movement after approximately two hours (Figure 5F). A summary of the phenotypic characteristics of each strain is provided in Table 3.

**Figure 4:**
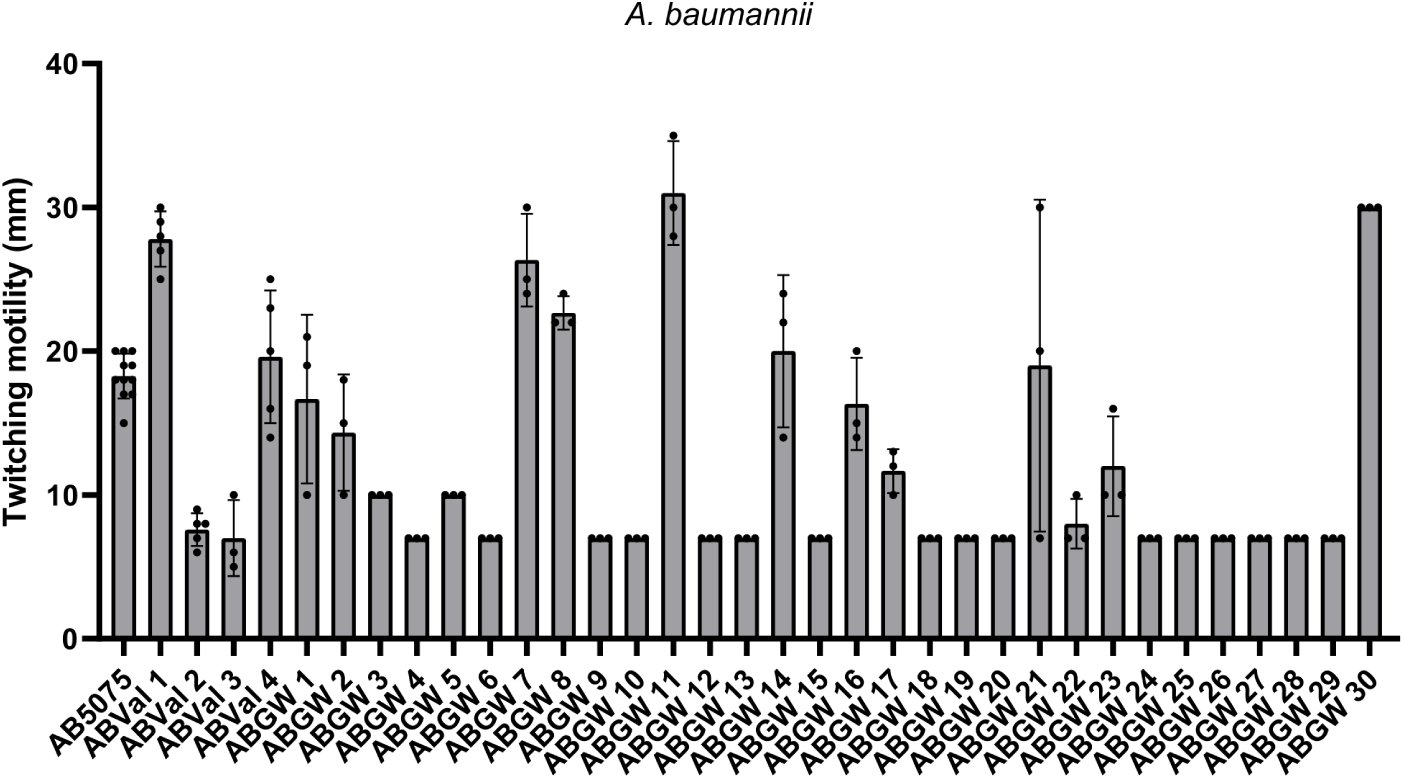
Comparative analysis of twitching motility among *A. baumannii* strains. Macroscopic twitching motility measured in millimeters (mm) after 24h for a set of *A. baumannii* strains, including AB5075, hospital strains, and environmental strains *(Note: ABVal 5 has been removed from the analyzed dataset)*. Three independent replicates were performed for each strain. Only strains displaying displacements greater than 10 mm exhibit twitching motility, whereas strains with movements below this threshold are not considered to perform twitching motility.

**Figure 5.**
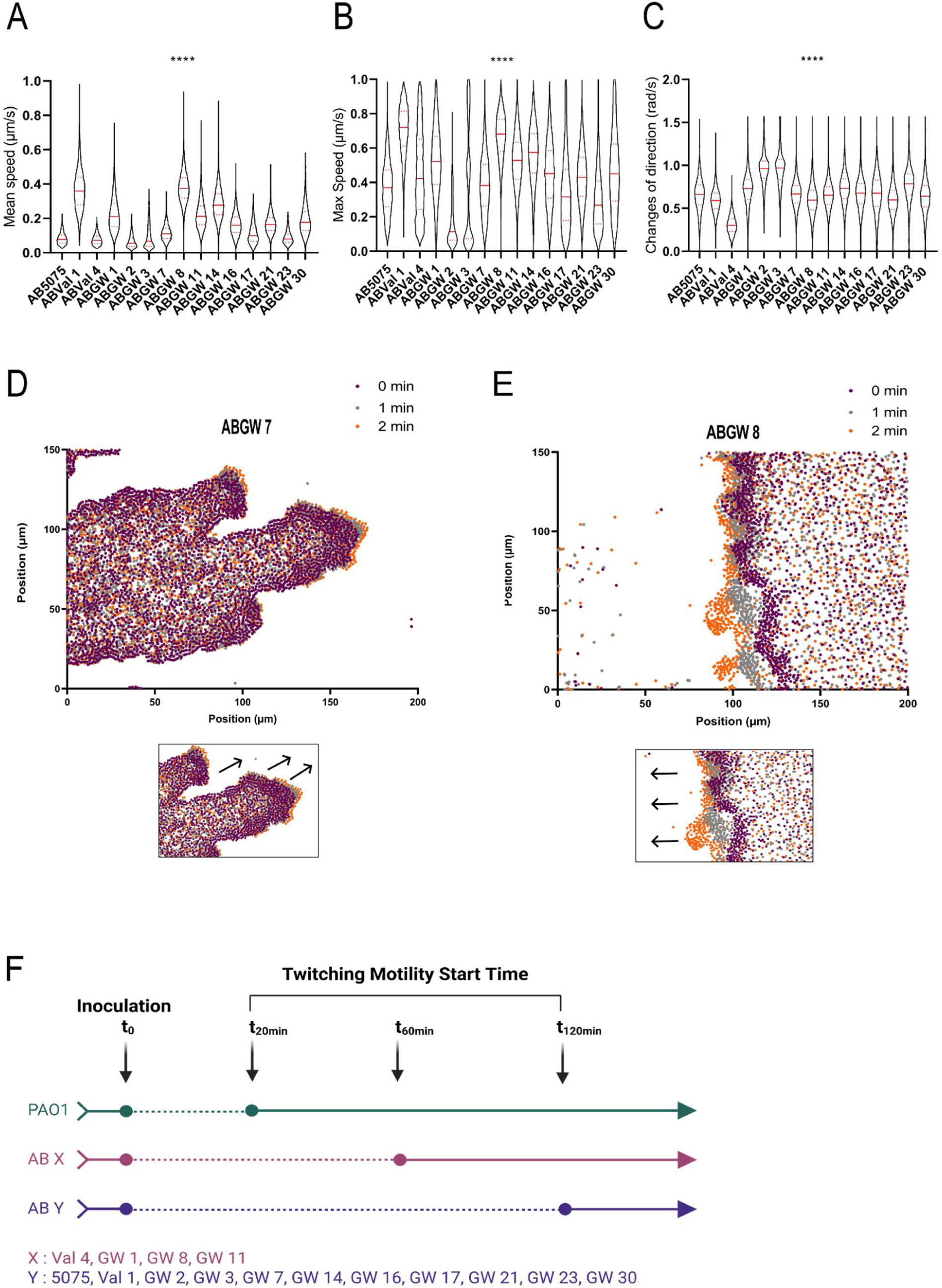
Microscopic characterization of twitching motility and collective migration patterns in *A. baumannii* strains. Microscopic analysis was performed on *A. baumannii* strains exhibiting macroscopic twitching motility. Quantitative analysis of bacterial movement two hours after inoculation included (A) average speed (µm/s), (B) maximum speed (µm/s), and (C) directional changes (rad/s). These parameters were extracted from time-lapse video microscopy and measured in three independent experiments. Representative trajectories illustrating distinct collective migration behaviors are shown in (D) and (E). Bacterial positions are displayed at three time points: T = 0 min (purple), T = 1 min (gray), and T = 2 min (orange), recorded 5 h after inoculation. (D) Strain ABGW 7 forms elongated migration arms reminiscent of the tentacle-like structures described for *P. aeruginosa* PAO1, as highlighted in the magnified panel. Black arrows indicate the principal axes of arm extension. (E) In contrast, strain ABGW 8 displays a coordinated and homogeneous migration front similar to that observed for strain AB5075, with black arrows indicating the overall direction of front progression. (F) Schematic representation comparing the onset of twitching motility after inoculation among the different *A. baumannii* strains, illustrating the diversity in initiation timing of surface-associated movement. Statistical significance was determined using the Kruskal-Wallis test following two-way analysis of variance (****, p ≤ 0.0001; *, p ≤ 0.05; ns, p > 0.05).

**Table 1.** Bacterial strains and plasmids used in this study

| Bacterial species | Strain or plasmid | Source or reference |
| --- | --- | --- |
| *Pseudomonas aeruginosa* | PAO1 | (Holloway and Morgan, 1986) |
|  | PAO1 *pilA::Tn5 ISlacZ/hah* | PAO1 transposon mutant library - Manoil Lab UW genome Sciences |
|  | PAO1 *pilT::Tn5 ISlacZ/hah* |  |
| *Acinetobacter baumannii* | AB5075 | (Jacobs et al., 2014) |
|  | AB5075 *pilA::T26* | AB5075 transposon mutant library - Manoil Lab UW genome Sciences |
|  | AB5075 *pilT::T26* |  |
| | AB5075 $\Delta fimT$ | This study |
|  | ABVal 1 | (Dessenne et al., 2024) |
|  | ABVal 2 |  |
|  | ABVal 3 |  |
|  | ABVal 4 |  |
|  | ABGW 1 (13-150-1C) | (Wilharm et al., 2026) |
|  | ABGW 2 (13-156-2C) |  |
|  | ABGW 3 (13-268-2C) |  |
|  | ABGW 4 (13-278-3C) |  |
|  | ABGW 5 (13-284-2C) |  |
|  | ABGW 6 (13-291-1C) |  |
|  | ABGW 7 (13-658C) |  |
|  | ABGW 8 (14-1PW1) |  |
|  | ABGW 9 (14-2P2) |  |
|  | ABGW 10 (14-2PW1) |  |
|  | ABGW 11 (14-73D4) |  |
|  | ABGW 12 (14-9D1) |  |
|  | ABGW 13 (14-Fe-1) |  |
|  | ABGW 14 (14-Spoe1-1) |  |
|  | ABGW 15 (14-Spoe3-1) |  |
|  | ABGW 16 (15-Entamr1) |  |
|  | ABGW 17 (15-Geschum2) |  |
|  | ABGW 18 (15-Lo_HL620-1) |  |
|  | ABGW 19 (15-LoFedIst2-1) |  |
|  | ABGW 20 (15-LoFedIst4-1) |  |
|  | ABGW 21 (15-LoHL619-1) |  |
|  | ABGW 22 (15-NCu29D-2) |  |
|  | ABGW 23 (15-NCu32R24h) |  |
|  | ABGW 24 (15-TF22) |  |
|  | ABGW 25 (16-Reih3-Ei-1) |  |
|  | ABGW 26 (16-SUI21-2) |  |
|  | ABGW 27 (16W1E1) |  |
|  | ABGW 28 (17-HoPe_P1-2) |  |
|  | ABGW 29 (17-HoPe_S1-2) |  |
|  | ABGW 30 (17-Hope_S16-5) |  |
| *Escherichia coli* | Top 10'F | Invitrogen |
| | S17 $\lambda$ pir | (Simon et al., 1983) |
| Plasmids | pASG-1 | (Godeux et al., 2018) |
|  | pASG-*fimT* | This study |
|  | pGP704-Sac18-amp, Suicide plasmid, oriR6K sacB ampR | (Metzger et al., 2019) |
|  | pGP704-SacB-Apr | This study |
| | pGP704-SacB-Apr-$\Delta fimT$ | This study |

**Table 2.**
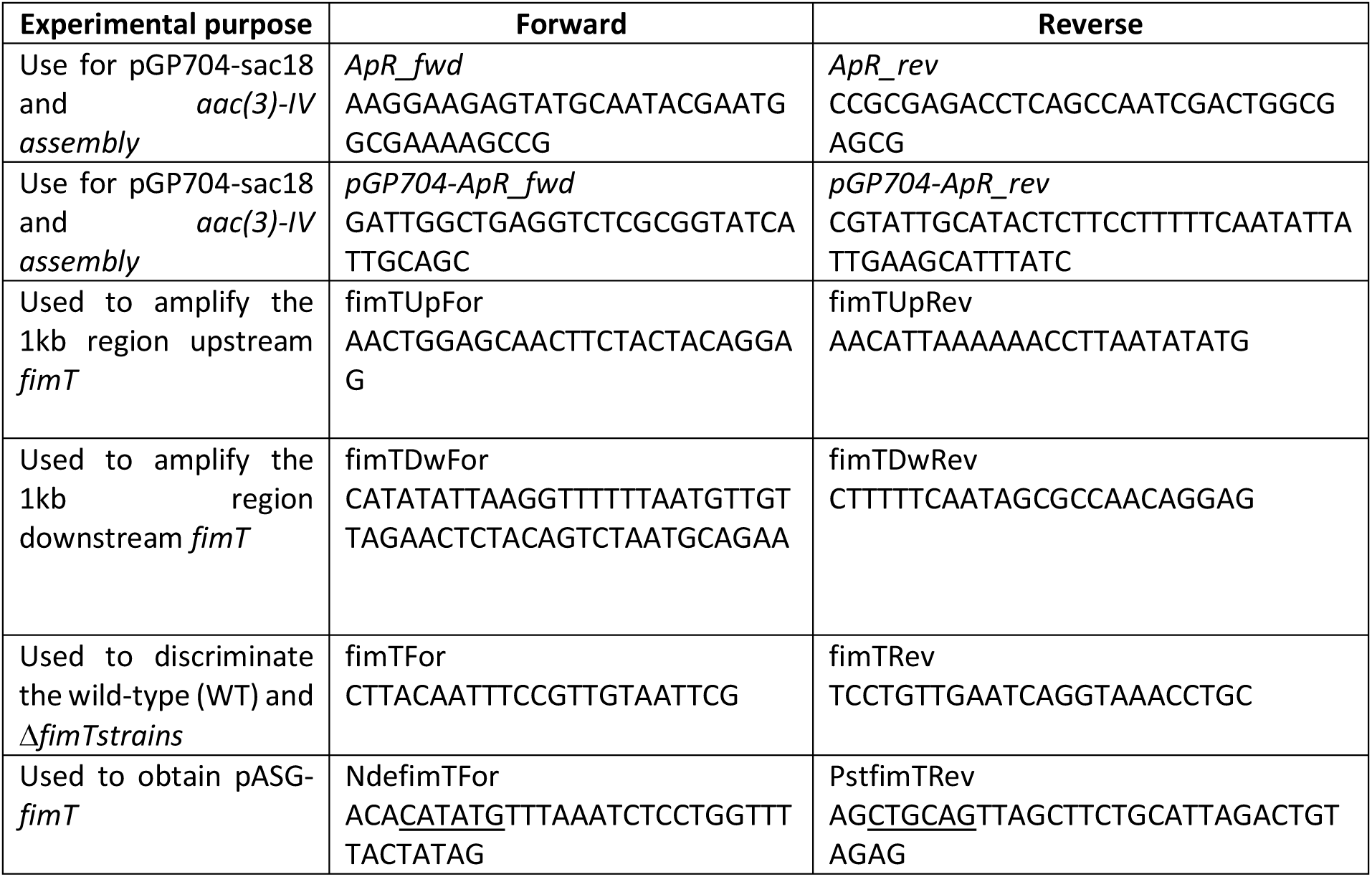
Primers used in this study.

**Table 3:** Twitching motility characteristics of PAO1 and 14 *A. baumannii* strains. Bacterial movement indicates the pattern of surface translocation: HF = Homogeneous Front, with smooth, uniform radial expansion; RLF = Raft-Like Front, with patchy, loosely aggregated clusters; R = Rafts, as observed for PAO1. Start twitching motility denotes the time required for detectable movement: EOM = Early-Onset Motility (∼1 h post-inoculation), DOM = Delayed-Onset Motility (∼2 h post-inoculation), or explicit times for PAO1.

| Strains | Type of motility | Start of motility |
| --- | --- | --- |
| PAO1 | R | 20 min |
| AB5075 | HF | DOM |
| ABVal 1 | RLF | DOM |
| ABVal 4 | RLF | EOM |
| ABGW 1 | RLF | EOM |
| ABGW 2 | RLF | DOM |
| ABGW 3 | RLF | DOM |
| ABGW 7 | RLF | DOM |
| ABGW 8 | HF | EOM |
| ABGW 11 | HF | EOM |
| ABGW 14 | RLF | DOM |
| ABGW 16 | RLF | DOM |
| ABGW 17 | RLF | DOM |
| ABGW 21 | HF | DOM |
| ABGW 23 | RLF | DOM |
| ABGW 30 | RLF | DOM |

### FimT contributes to twitching motility but does not fully explain motility heterogeneity

Because 21 of the 35 strains examined were non-motile, we investigated whether sequence variation in the minor pilin FimT could account for the observed heterogeneity. Previous work identified several substitutions potentially associated with impaired twitching motility, including the I9T mutation affecting the prepilin processing motif and the recurrent R91H substitution (Iruegas et al., 2023). Comparison of FimT amino acid sequences across the strain collection revealed fifteen substitutions (Supplementary Fig. S1). Two substitutions (T13A and L150F) were found exclusively in motile strains. Four substitutions (D71N, V83L, R91H, and S131N) were detected in non-motile isolates but were also present in motile strains, indicating that they could not independently explain the loss of motility. Only three substitutions, I9T, N108D, and N152K, were unique to non-motile strains. To evaluate the contribution of FimT to twitching motility, a Δ*fimT* mutant was generated in the AB5075 background. Deletion of *fimT* completely abolished twitching motility (Figure 6A, AB Δ*fimT*pASG), whereas complementation with the AB5075 *fimT* allele restored movement around to 60% (Figure 6B, AB5075 Δ*fimT*pASG-*fimT*), confirming the essential role of FimT in T4aP-dependent motility. We next assessed whether the non-motile GW strains could be rescued by the AB5075 *fimT* allele. Complementation restored twitching in ABGW6 (I9T) and ABGW27 (N152K) to approximately 80% and 70% of WT levels, respectively, but failed to restore motility in ABGW5 (N108D) (Figure 6A), suggesting that additional factors contribute to the non-motile phenotype of this strain. To directly evaluate the functional impact of individual FimT variants, the corresponding alleles were expressed separately in the AB5075 Δ*fimT* background (Figure 6B). The I9T and N108D variants restore substantial motility, with approximately 75% and 65% of WT motility, respectively. In contrast, N152K completely abolished twitching, whereas R91H and S131N produced intermediate phenotypes with approximately 50% reductions in motility. To investigate the structural basis of these phenotypes, predicted models of AB5075 FimT and the variants identified in ABGW5, ABGW6, and ABGW27 were compared. Structural superposition revealed highly conserved global folds (Figure 6C), with RMSD values below 0.7 Å for all pairwise comparisons, indicating that none of the substitutions caused major conformational changes (Supplementary Tables 1A and B). Consistent with their high evolutionary conservation, residues N108 and N152 appear to be under stronger functional constraints than the more variable residue I9. While the I9T and N108D substitutions induced only subtle local structural changes, the N152K substitution caused a more pronounced rearrangement around residues 151–152. Electrostatic surface analysis further showed that I9T did not alter the local charge distribution (Supplementary Fig. S2A,B), whereas N108D generated a localized increase in negative surface potential (Supplementary Fig. S2A’,C) and N152K produced a marked positive electrostatic patch, further enhanced by the local structural rearrangement (Supplementary Fig. S2A’’,D). Thus, although all variants preserve the overall FimT fold, the N108D and especially the N152K substitutions substantially remodel the local electrostatic environment, providing a plausible structural explanation for their reduced ability to support twitching motility.

**Figure 6.**
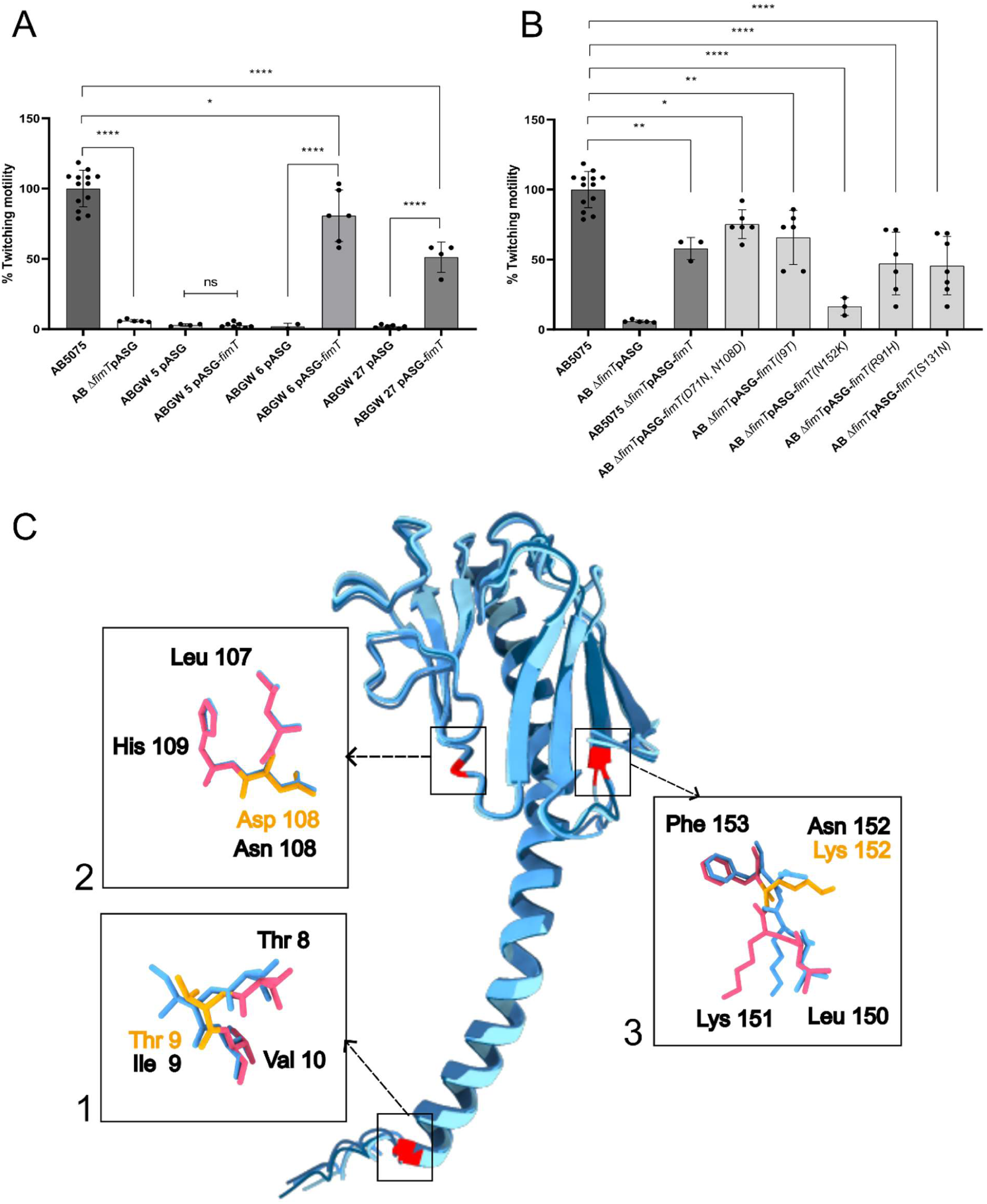
Functional and structural analysis of natural FimT variants associated with twitching motility in *A. baumannii*. Macroscopic twitching motility was quantified after 48 h of incubation. (A) Functional evaluation of the WT *fimT* allele, the Δ*fimT* deletion mutant, and the indicated *A. baumannii* GW isolates (ABGW 5, ABGW 6, and ABGW 27), together with genetic complementation of the Δ*fimT* mutant and the GW strains using the WT *fimT* allele from strain AB5075 expressed from the pASG vector. (B) Complementation of the non-motile AB5075 Δ*fimT* mutant with plasmids expressing the different *fimT* alleles amplified from the respective ABGW strains. Non-motile controls (Δ*fimT* mutant and empty pASG vector) were set to zero. Strains exhibiting a twitching zone greater than 40% of the WT level were considered motile. (C) Structural comparison of FimT variants. Global superposition of the AB5075 FimT structure with the variants from ABGW 5, ABGW 6, and ABGW 27. Residues from the WT AB5075 protein are shown in blue, whereas those from the GW variants are shown in pink. The substituted amino acid is highlighted in orange. Enlarged views illustrate the local structural environments surrounding the I9T, N108D, and N152K substitutions. Statistical significance was determined using Tukey’s post hoc test following one-way analysis of variance (ANOVA) (****, *p* ≤ 0.0001).

## Discussion

Our results reveal that twitching motility in *A. baumannii* is more diverse than previously appreciated. Beyond differences in motility speed, we identified substantial variation in the timing of twitching initiation and in the collective organization of migrating populations. These observations suggest that the regulation and deployment of T4P, rather than their mere presence, play a central role in determining surface colonization strategies in this species. More than two decades ago, Mattick proposed that twitching motility should be viewed as a dynamic multicellular behavior whose manifestation depends on both species-specific regulation and environmental conditions rather than as a single conserved phenotype (Mattick, 2002). While this conceptual framework was largely established from studies of *P. aeruginosa*, experimental evidence from other bacterial species has remained limited. One of the most striking differences observed between *A. baumannii* and *P. aeruginosa* was the timing of twitching initiation. Under identical experimental conditions, PAO1 initiated twitching within approximately 15 minutes after inoculation, whereas AB5075 remained non-motile for nearly two hours before surface translocation became detectable. Importantly, this delayed onset was not restricted to AB5075 but was observed in the majority of motile *A. baumannii* strains, suggesting that delayed twitching may represent a common feature of the species. The mechanisms underlying this latency remain unclear. Because T4P-associated genes are expressed during growth and have been shown to be induced under host-relevant conditions (Jacobs et al., 2014; Murray et al., 2017; Vesel and Blokesch, 2021), the delay is unlikely to reflect a simple absence of transcription. Instead, it may arise from post-transcriptional regulation, delayed pilus assembly, activation of extension-retraction cycles, or surface-sensing mechanisms that must be engaged before productive motility can occur. Similar regulatory checkpoints have been described in *P. aeruginosa*, where surface contact triggers complex signaling pathways involving PilY1, cyclic-di-GMP, and mechanosensory systems that coordinate the transition from planktonic to surface-associated lifestyles (Ellison et al., 2022; Laventie and Jenal, 2020). Whether comparable pathways regulate twitching onset in *A. baumannii* remains unknown in our knowledge. The observation that some strains exhibited EOM, whereas others displayed DOM, further suggests that substantial variation exists in the regulatory networks controlling T4P deployment. Future studies directly quantifying pili abundance and localization over time will be required to determine whether delayed motility reflects delayed pilus production, delayed activation of already assembled pili, or density-dependent collective effects. Importantly, the variability observed in twitching phenotypes is unlikely to arise solely from biological differences between strains. The pronounced effect of agar composition observed in this study suggests that experimental conditions may also contribute to discrepancies in twitching prevalence reported across *A. baumannii* collections. Previous studies have shown that surface hydrophilicity, medium composition, and agar concentration can strongly influence T4P-dependent motility phenotypes, supporting the need for standardized assay conditions when comparing strains or studies (Biswas et al., 2019; O’Hara et al., 2024).

Beyond differences in twitching onset, motile *A. baumannii* strains displayed two distinct modes of collective organization. Some strains formed homogeneous fronts (HF) characterized by smooth radial expansion, whereas others exhibited raft-like fronts (RLF) composed of loosely associated multicellular clusters. These phenotypes reveal that collective migration strategies vary substantially within the species. Interestingly, neither organization resembled the highly structured finger-like projections formed by *P. aeruginosa* PAO1. In *P. aeruginosa*, twitching motility generates multicellular rafts whose behavior emerges from physical interactions between cells and local packing constraints rather than from the behavior of individual bacteria alone (Meacock et al., 2021). Under these conditions, collective performance can become partially uncoupled from individual cell speed, allowing slower but more coordinated populations to colonize surfaces more efficiently than faster but less organized groups. Our observations suggest that similar emergent phenomena may operate in *A. baumannii*. Although strains differed in mean speed, maximum speed, and angular velocity, these parameters alone did not predict colony-front architecture. Instead, HF and RLF phenotypes likely reflect differences in how cells coordinate T4P-mediated interactions at the population level. Variations in local cell density, pilus-pilus interactions, adhesion strength, or reorientation dynamics may all contribute to the emergence of distinct collective behaviors. Such diversity may have important ecological consequences. Different collective migration strategies could influence how rapidly strains colonize surfaces, compete within polymicrobial communities, or initiate biofilm formation. The broad distribution of HF, RLF, EOM, and DOM phenotypes across genetically diverse isolates suggests that twitching behavior is not a single conserved trait but rather a flexible adaptation shaped by ecological context.

Although approximately 40% of the strains examined displayed twitching motility, the remaining 60% were non-motile despite carrying many of the genes associated with T4P biogenesis. This finding is consistent with previous reports indicating that twitching motility is highly variable among *A. baumannii* isolates and cannot be predicted solely from the presence of T4P-associated genes (Al-Shamiri et al., 2021; Corral et al., 2021; Eijkelkamp et al., 2011). To investigate whether variation in the minor pilin FimT contributed to this diversity, we combined sequence analyses, complementation experiments, and structural modeling. Our data clearly demonstrate that FimT is required for twitching motility in AB5075, as deletion of the gene abolished motility and complementation restored the phenotype. This contrasts with observations in *Acinetobacter baylyi* and *Pseudomonas* species, where FimT appears dispensable for twitching motility (Alm and Mattick, 1996; Leong et al., 2017), suggesting that the contribution of this minor pilin varies across species. Nevertheless, FimT variation alone could not explain the broad diversity of twitching phenotypes observed in our collection. Most substitutions identified among non-motile isolates were also present in motile strains, and complementation experiments demonstrated that several naturally occurring variants retained substantial activity. Only the N152K substitution produced a severe loss-of-function phenotype, completely abolishing twitching when expressed in a Δ*fimT* background. Structural analyses indicated that this mutation does not disrupt the global fold of FimT but instead alters local electrostatic properties within the conserved C-terminal region. This region has previously been implicated in DNA binding during natural transformation (Braus et al., 2022), suggesting that subtle modifications of surface charge may influence interactions with partner proteins or pilus assembly intermediates. These findings indicate that FimT contributes to T4P function and twitching motility in *A. baumannii*, but it is unlikely to be the primary determinant of motility heterogeneity at the population level. The persistence of non-motile phenotypes despite functional FimT variants strongly suggests that additional structural, regulatory, or physiological factors remain to be identified. Future work combining genomics, transcriptomics, and direct measurements of pilus production will be required to determine how these factors interact to shape the remarkable diversity of twitching behaviors observed across *A. baumannii* strains. Overall, our work revisits the conceptual framework established by Mattick (Mattick, 2002) for T4P-dependent twitching motility using *A. baumannii*, the bacterial genus in which twitching was originally described by Lautrop (Lautrop, 1961). Rather than identifying a single conserved mode of surface translocation, we reveal that twitching motility encompasses diverse temporal programs, collective migration strategies, and genetic determinants. These findings broaden the current view of T4P-mediated motility and provide a foundation for investigating how natural variation in multicellular behavior contributes to environmental adaptation, host colonization, and virulence.

---

**Supplementary Figure S1:**
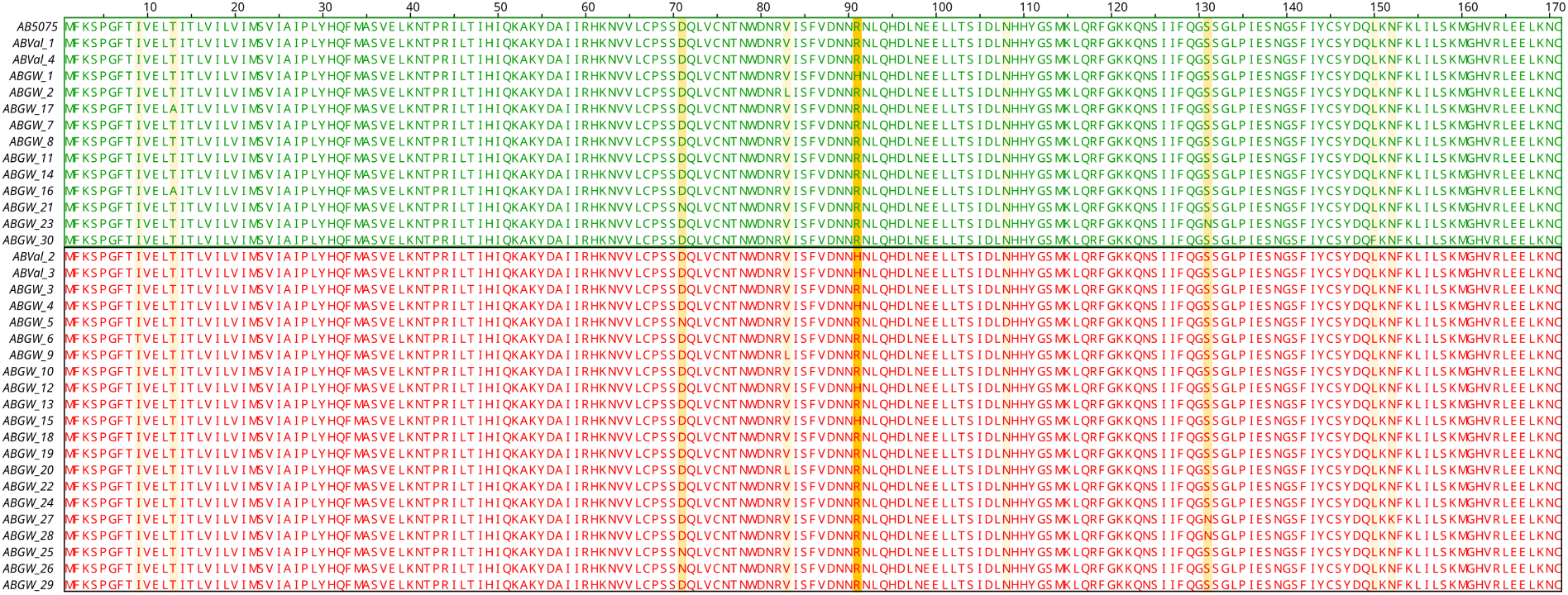
Sequence alignment of FimT across *A. baumannii* strains and correlation with twitching phenotype. Multiple sequence alignment of GspH protein sequences from different *A. baumannii* strains. Amino acid differences relative to AB5075 are highlighted to illustrate sequence variability among strains. Strains exhibiting twitching motility are indicated in green, whereas non-twitching strains are indicated in red.

**Supp. Figure S2:**
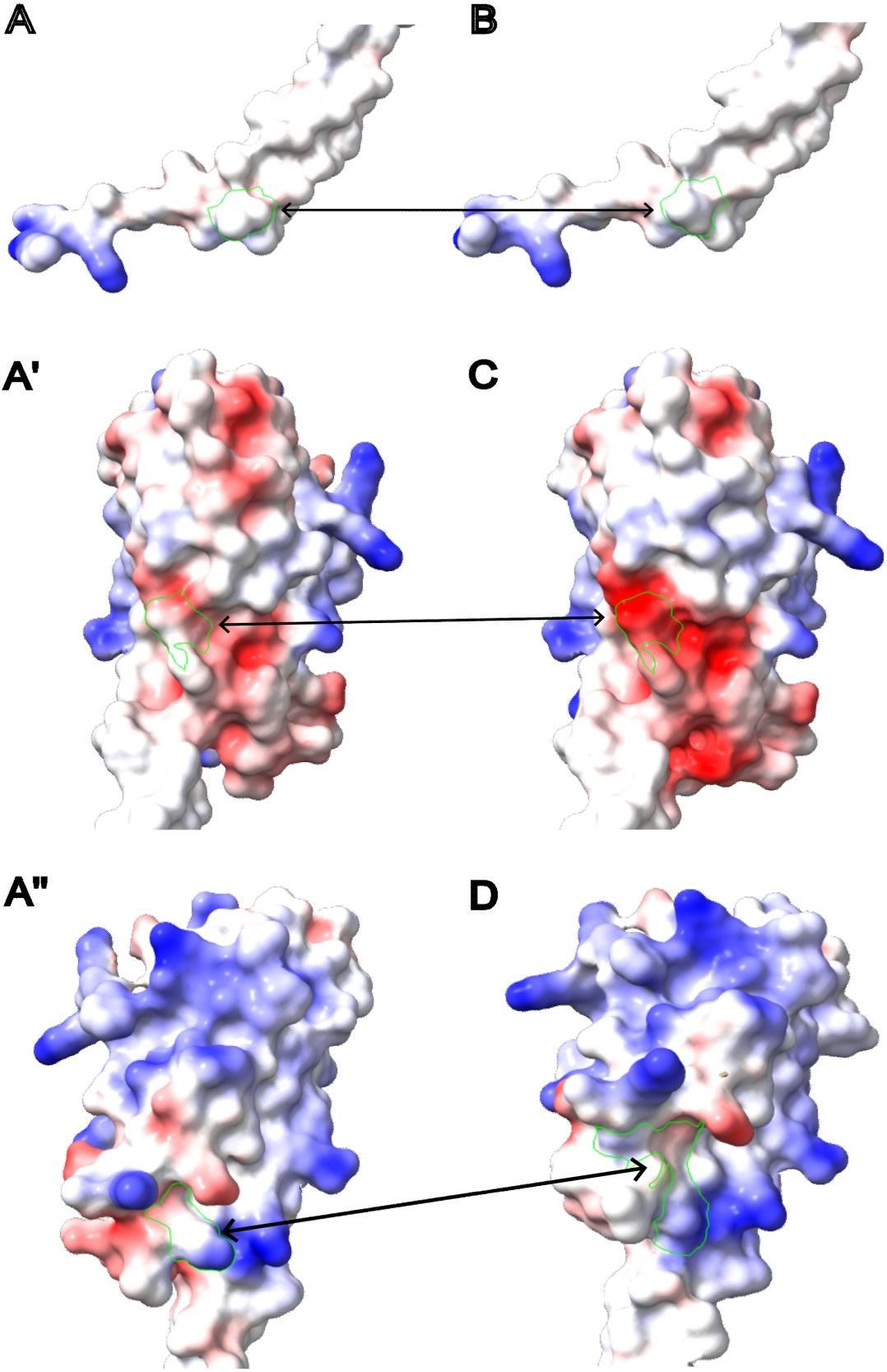
Comparative electrostatic potential surface analysis of FimT variants. Local electrostatic surface potentials were calculated using Coulombic surface coloring in ChimeraX, where blue, red, and white represent positively charged, negatively charged, and neutral regions, respectively. (A-A’’) Electrostatic profiles of the WT AB5075 reference strain, turned and oriented specifically toward the corresponding residue position to be compared: position 9 (A), position 108 (A’), and position 151-152 (A’’). (B) The I9T variant (strain ABGW 6) displays an identical electrostatic profile to its corresponding WT orientation (A), demonstrating that this substitution has no impact on the local surface charge distribution. (C) The N108D variant (strain ABGW 5) results in a highly significant localized increase in negative charge (expanded red area) compared to (A’), driven by the substitution of a neutral asparagine with a negatively charged aspartic acid. (D) The N152K variant (strain ABGW 27) induces a strong local shift toward a positive electrostatic potential (blue area) compared to (A’’), resulting from both the introduction of a positively charged lysine residue and its associated local structural rearrangement.

**Supp. Table 1:** Structural comparison and alignment metrics of FimT variants. (A) Global RMSD values calculated from ChimeraX superpositions between the WT *A. baumannii* AB5075 FimT protein and the indicated mutant variants, confirming a highly conserved overall global fold. (B) Local, residue-specific structural metrics surrounding the mutated sites, including local RMSD values, Cα distances, solvent-accessible surface area (SASA), and electrostatic or side-chain-specific measurements. These local metrics highlight distinct residue-specific structural perturbations, most notably the strong conformational disruption around residue 151 induced by the N152K variant. “-” indicates not measured or not applicable.

| Comparison / Residue | RMSD pruned (Å) | RMSD all pairs (Å) |
| --- | --- | --- |
| AB5075 vs ABGW 5 | 0.6 | 0.67 |
| AB5075 vs ABGW 6 | 0.516 | 0.739 |
| AB5075 vs ABGW 27 | 0.585 | 0.687 |

B
| Strains | Mutation | Comparison / Residue | RMSD pruned (Å) | Distance CA (Å) | RMSD N, CA, C, O (Å) | RMSD @NZ (Å) | SASA WT (AB5075) | SASA mutant | Comment |
| --- | --- | --- | --- | --- | --- | --- | --- | --- | --- |
| ABGW 5 | N108D | Residue 107 (L) | 0.335 | 0.352 | 0.351 | - | 292.5 | 292.5 | Local environment around mutation |
|  |  | Residue 108 | 2.65 | 0.309 | 0.295 | - | 263.17 | 256.89 | Mutated residue |
|  |  | Residue 109 (H) | 0.265 | 0.231 | 0.234 | - | 297.97 | 298.01 | Local environment around mutation |
| ABGW 6 | I9T | Residue 8 (T) | 1.074 | 1.131 | 1.103 | - | 248.08 | 248.09 | Local environment around mutation |
|  |  | Residue 9 | - | 0.983 | 0.877 | - | 287.33 | 251.28 | Mutated residue |
|  |  | Residue 10 (V) | 0.579 | 0.588 | 0.611 | - | 262.05 | 262.05 | Local environment around mutation |
| ABGW 27 | N152K | Residue 151 (K) | 3.506 | 1.863 | 1.45 | 6.424 | 314.49 | 313.98 | Most affected local residue |
|  |  | Residue 152 | - | 0.996 | 1.13 | - | 261.34 | 310.66 | Mutated residue |
|  |  | Residue 153 (F) | 0.58 | 0.704 | 0.676 | - | 310.95 | 310.98 | Local environment around mutation |

## Author contributions

CD: formal analysis, investigating, methodology, validation, visualization. AH: formal analysis and investigating. AC: formal analysis and investigating. CS: methodology. DD: formal analysis and investigating. OV: conceptualization, resources, supervision, writing, review and editing. YR: conceptualization, funding acquisition, investigating, methodology, project administration, supervision, writing original draft.

## Acknowledgements

We thank the Plateformes Lilloises en Biologie et Santé (PLBS) - UAR 2014 - US 41, for providing access to instrumentation and for their valuable technical support. We are grateful to Gottfried Wilharm for generously providing 30 *A. baumannii* strains. We also thank Xavier Charpentier for the pASG-1 plasmid, and Mélanie Blokesch for her advice on generating *A. baumannii* mutants and for kindly providing the Escherichia coli S17 λpir strain.

